# Unbiased definition of a shared T-cell receptor motif enables population-based studies of tuberculosis

**DOI:** 10.1101/123174

**Authors:** W. S. DeWitt, K. K. Quan, D. Wilburn, A. Sherwood, M. Vignali, S. C. De Rosa, C. L. Day, T. J. Scriba, H. S. Robins, W. Swanson, R. O. Emerson, C. Seshadri

## Abstract

Peptide-specific T cells that are restricted by highly polymorphic major histocompatibility complex (MHC) proteins express diverse T-cell receptors (TCRs) that are rarely shared among unrelated individuals. T-cells can also recognize bacterial lipid antigens that bind the relatively non-polymorphic CD1 family of proteins. However, genetic variation in human CD1 genes and TCR diversity expressed by CD1-restricted T-cells have not been quantitatively determined. Here, we show that CD1B is nearly nucleotide-identical across all five continental ancestry groups, providing evidence for purifying selection during human evolution. We used CD1B tetramers loaded with a mycobacterial glycolipid antigen to isolate T-cells from four genetically unrelated South African adults and cataloged thousands of TCRs from *in-vitro* expanded T-cells using immunosequencing. We identified highly conserved motifs that were co-expressed as a functional heterodimer and significantly enriched among tetramer-positive T-cells sorted directly from peripheral blood. Finally, we show that frequencies of these TCR motifs are increased in the blood of patients with active tuberculosis compared to uninfected controls, a finding that is confirmed by ex-vivo frequencies of tetramer-positive T-cells determined by flow cytometry. These data provide a framework for unbiased definition of TCRs targeting lipid antigens, which can be tested for clinical associations independently of host genetic background.

**Brief Summary:** We used human genetics and immunosequencing to define a shared T-cell receptor motif that is specific for a mycobacterial lipid antigen and associated with tuberculosis independently of host genetic background.

## INTRODUCTION

Classically, T cells recognize peptide antigens when bound to major histocompatibility complex (MHC) molecules (1). The six MHC class I and class II genes are among the most diverse in the human genome. MHC diversity provides a population advantage against pathogens by reducing the probability that a mutation will evade detection by the adaptive immune system. This leads to a selective preference for MHC heterozygosity (2). T cells express an antigen-specific T cell receptor (TCR), which enables recognition of peptide antigen only when it is bound to MHC. A consequence of MHC allelic diversity is that each host must develop a personalized set of T cells able to recognize peptide-based antigens produced by any pathogen. Even among identical twins, the stochastic nature of TCR formation results in largely distinct naïve T cell repertoires, although increased sharing is observed among antigen-experienced T cells (3). Thus, the collection of all T cells (repertoire) between two genetically unrelated individuals rarely overlaps.

The TCR is a heterodimer consisting of an α and β chain that is generated by somatic recombination of germline-encoded segments. Further junctional diversity is provided by nucleotide additions and deletions, increasing the potential diversity to nearly one trillion unique sequences (4, 5). However, the actual number of unique T cells (clonotypes) in peripheral blood is estimated at 3-4 million, a substantially lower number because of the processes of positive and negative selection that occur in the thymus and because of the finite number of T cells present in a single individual at a specific point in time (6). Among genetically unrelated individuals that share a dominant MHC allele, a minority of T cells recognizing a viral peptide antigen share a common TCR-β sequence (7). These examples demonstrate the challenge for generalizing information derived from TCRs without first conditioning on genetic similarity.

Notably, human T cells have evolved mechanisms independent of MHC to facilitate the recognition of non-peptide antigens. Such antigens include bacterial lipids, which bind the CD1 family of antigen-presenting molecules (8). There are five CD1 proteins in humans (CD1A, CD1B, CD1C, CD1D, and CD1E) capable of processing and presenting lipid antigens to T cells. Rapid evolution is common among immunity-associated genes, and MHC class I molecules are frequently under strong directional selection (9). By contrast, CD1 genes do not display the level of polymorphism inherent to MHC, though the actual levels of human genetic variation have not been quantitatively determined. CD1 gene conservation may result in CD1-restricted T cells with limited TCR diversity. For example, a number of studies have documented the presence of invariant NK T (iNKT) cells in the blood of unrelated subjects from different populations (10-12). These cells are activated by glycolipid antigens bound to CD1D, and canonically express a TCR-α consisting of a germline rearrangement including TRAV10-1 and TRAJ18-1 gene segments in humans (13, 14). Similarly, germline-encoded mycolyl lipid-reactive (GEM) T cells are activated by mycobacterial glycolipids bound to CD1B and express a TCR-α consisting of TRAV1-2 and TRAJ9-1 gene segments (15). Both iNKT and GEM cells express TCR-β with biased gene segment usage, thus permitting the recognition of multiple antigens.

Another class of T cells that are shared at the population level are mucosal-associated invariant T (MAIT) cells, which are activated by bacterial metabolites of vitamin B that bind the MR1 antigen-presenting molecule (16). Like iNKT and GEM cells, MAIT cells express a semi-invariant TCR-α consisting of TRAV-1 and TRAJ33 gene segments in humans, and are detectable in the blood of unrelated individuals (14, 17). MAIT cells are activated in an MR1-dependent manner by M.tb-infected cells, though the M.tb-specific ligand(s) remain to be defined (18, 19). By contrast, a number of mycobacterial lipid antigens presented by CD1 proteins to T cells have already been discovered (20). However, it is not known whether these lipid antigens are recognized by antigen-specific T cells bearing TCRs that are shared across a population or whether these TCRs are associated with disease.

Our data reveal a remarkable conservation of CD1B across diverse populations, which is in stark contrast to the polymorphism that is inherent in MHC. We hypothesized that this structural constraint would enable a lipid-antigen specific T-cell response to be shared across genetically unrelated members of a population. By analyzing the TCRs of T cells specific for a mycobacterial lipid antigen, we identified a conserved T cell response that was detectable in the blood of multiple unrelated individuals. Using TCR immunosequencing and flow cytometry, we show that this lipid-specific T cell population is expanded during tuberculosis infection and disease. Our data reveal one example of an antigen-specific shared T cell repertoire that could be leveraged to develop molecular markers for diseases in which lipid antigens are targeted by T cells, such as tuberculosis, psoriasis, or leukemia.

## RESULTS

### Comparing genetic variation in Human HLA-A and CD1B

To compare the variation between classical and non-classical antigen presenting molecules, we examined DNA sequence diversity in a representative MHC Class I gene (HLA-A) and CD1 gene (CD1B) using available data from Phase 3 of the 1000 Genomes Project (21). This resource includes whole genome sequences from 2,504 individuals covering all five major continental ancestry groups, and thus it serves as a comprehensive resource for studying human genetic diversity. For each of these two loci, we quantified nucleotide diversity (π) and evidence of selection (Tajima’s D) at each position along the gene (22, 23). Within HLA-A, the median value of π was 0.015, with hotspots noted in exons 2 and 3, which are known to code for the peptide antigen-binding domains. We also observed a median value of 0.92 for Tajima’s D, suggesting balancing selection of multiple alleles at high and low frequencies at the population level (Figure 1A). We then examined the consequences of the observed nucleotide diversity by mapping these onto the crystal structure of HLA-A (24). This analysis highlights the distribution of missense, nonsense, and frameshift mutations that are particularly enriched among residues lining the peptide-binding groove (Figure 1B). These data are consistent with published studies of population diversity within MHC genes, and with our current understanding of how sequence diversity leads to diversity in ligand binding at the molecular level (25, 26). By contrast, CD1B was nearly invariant with a median π of 0.00014, and a median Tajima’s D value of −1.14, suggesting neutrality or weak purifying selection at the population level (Figure 1C). The tertiary structure of the lipid-binding domain of CD1B was completely conserved with only two synonymous polymorphisms outside of the antigen-binding groove (Figure 1D) (27). These data support purifying selection on CD1B during human evolution and stand in stark contrast to the immense diversity inherent within HLA-A and other MHC Class I genes.

**Figure 1.**
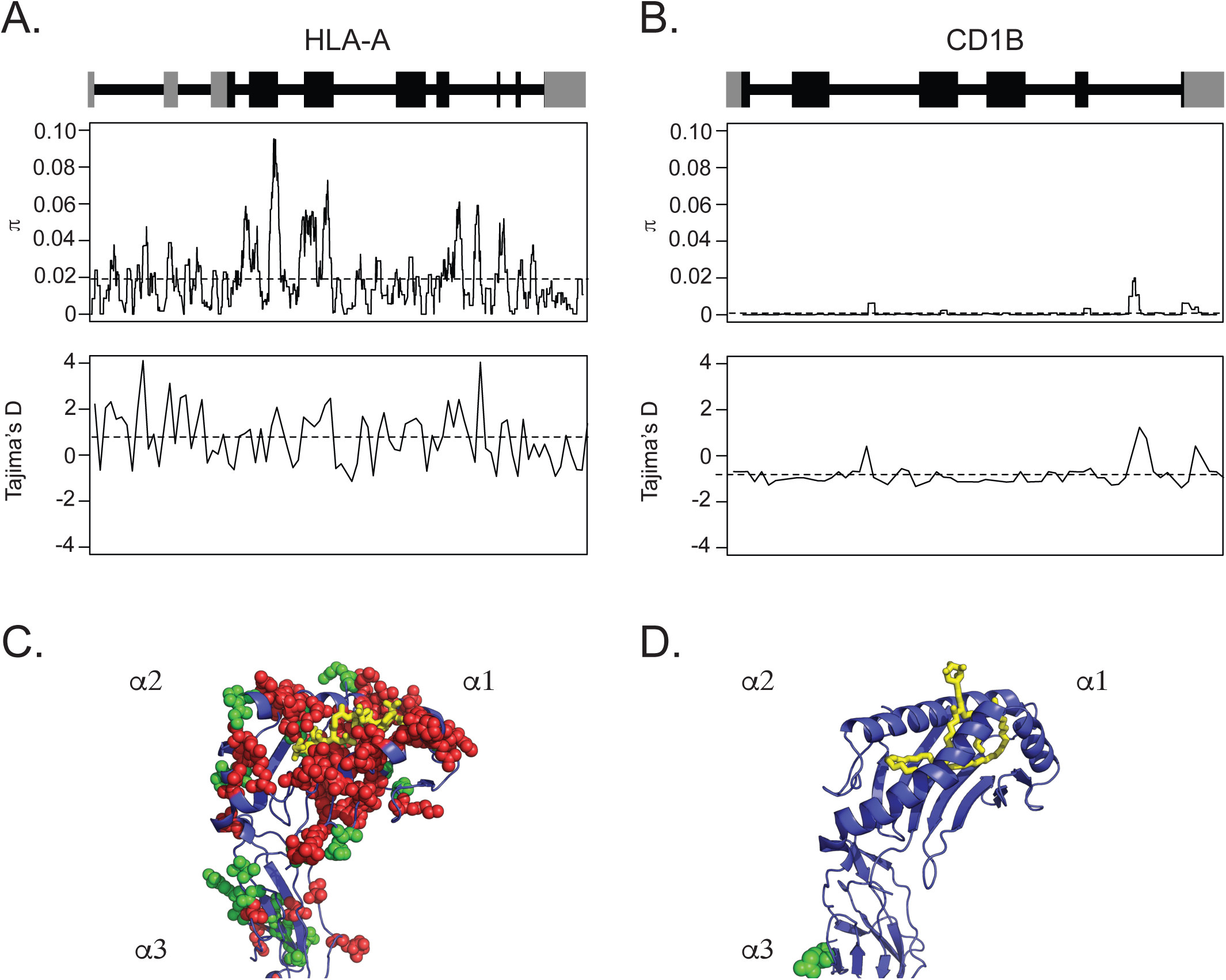
Human genetic variation in HLA-A and CD1B. The schematic of each gene structure, including introns (as lines) and exons (as blocks), with coding sequence indicated in black and untranslated regions in grey. Nucleotide diversity (π) and Tajima’s D are reported for (A) HLA-A2 or (B) CD1B with the average value across each gene denoted by a horizontal dashed line. Variation in protein coding sequence for (C) HLA-A2 or (D) CD1B is represented for SNPs with >1% minor allele frequency. Invariant residues are denoted in blue as a ribbon structure. Spheres denote polymorphic residues, with green representing synonymous substitutions and red representing non-synonymous substitutions. The ligand, either peptide or lipid, is shown in yellow.

### Derivation of GMM-specific T cell lines from healthy South African donors

The lack of allelic diversity within CD1B suggests that genetically unrelated individuals might generate a shared T-cell response to a given lipid antigen. To test this hypothesis directly, we used CD1B tetramers loaded with the immunodominant mycobacterial lipid glucose monomycolate (GMM) to isolate antigen-specific T cells from four healthy South African adults. Tetramer-positive cells were expanded *in-vitro* for four weeks to derive GMM-specific T cell lines (Figure 2A). We confirmed that GMM-specific T cells were primarily CD4+, that they were activated and produced IFN-γ in the presence of GMM and CD1b, and that they did not cross-react even with closely related antigens, such as mycolic acid (Figure 2B and data not shown). Finally, we confirmed that GMM-specific T cell lines were functional, expressing IFN-γ, TNF-α, IL-2, IL-17, and CD40L when stimulated though cognate interactions with antigen or independently of the T-cell receptor (Figure 2C and data not shown). These data confirm previously published data by our group and others showing that GMM-specific T cells with similar functional profiles are present in the blood of unrelated donors (28, 29).

**Figure 2.**
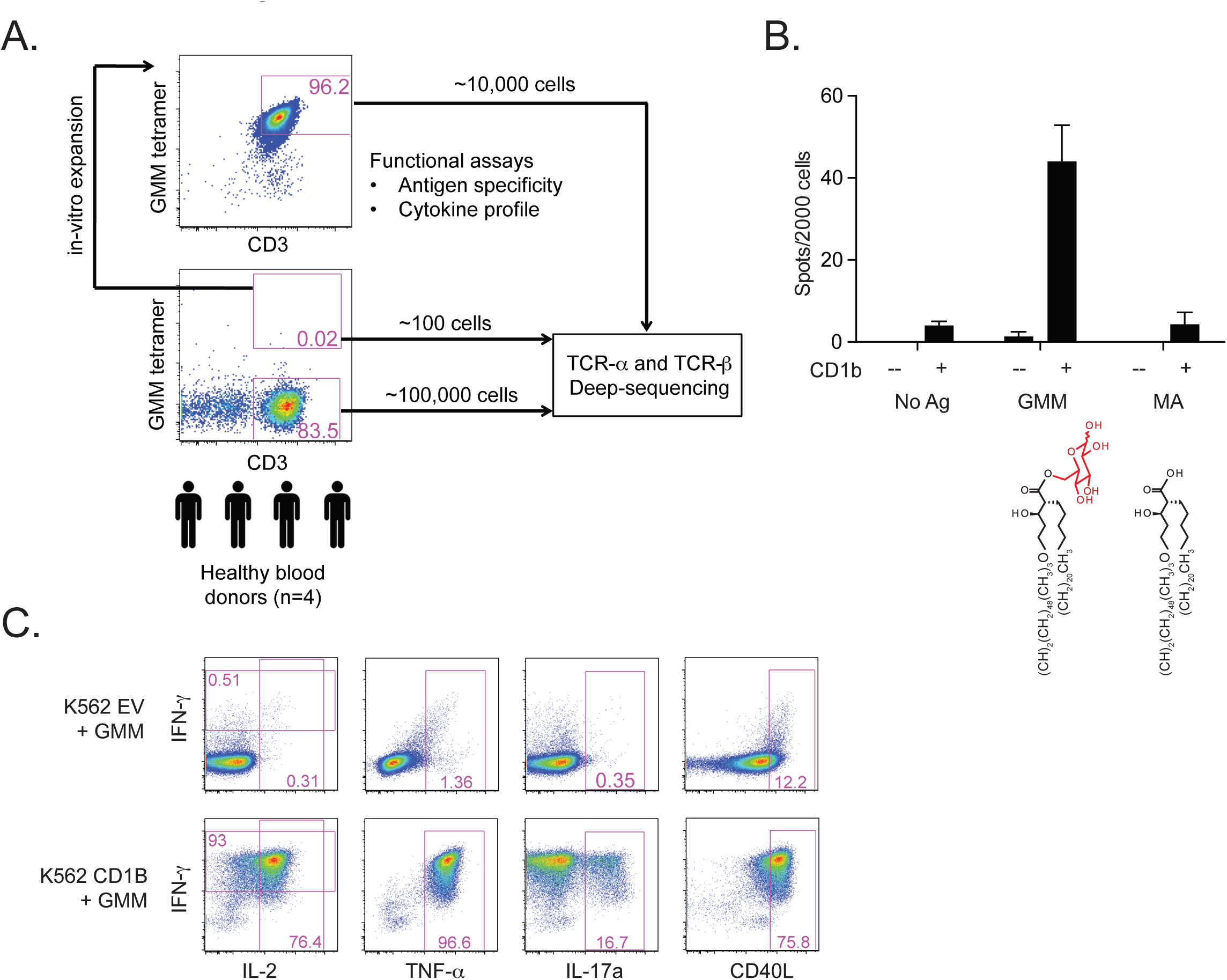
Study schema and characterization of T-cell lines. (A) We used GMM-loaded CD1B tetramers to stain and sort T cells from cryopreserved PBMCs derived from four healthy South African blood donors. These cells were either used to generate T-cell lines, or submitted directly for TCR immunosequencing. Tetramer-negative T cells as well as tetramer-positive T cells that were re-sorted from T cell lines were also submitted for TCR immunosequencing. (B) T-cell lines were incubated with either mock transfected or CD1B-transfected K562 cells as antigen-presenting cells in the presence of no antigen, GMM, or mycolic acid (MA). Partial structures of the two lipid antigens are shown. IFN-γ production by ELISPOT was quantified after overnight incubation. (C) Mock-transfected or CD1B-transfected K562 cells were loaded with GMM overnight and used to activate T cells for six hours prior to intracellular cytokine staining. Data in B and C are representative of three independent experiments.

### The diversity of T cells that bind CD1B-GMM tetramer

We then re-sorted those T cells that bound GMM-loaded CD1b tetramer with high avidity and exhaustively profiled the TCRs in both the tetramer-positive and tetramer-negative cell populations using high-throughput immunosequencing (Figure 2A)(6). The surprising diversity of V and J gene segments present in both TCR-α and TCR-β sequences from tetramer-positive cells suggest that a number of clonotypes (i.e., unique T-cells expressing a specific TCR-α and TCR-β) are able to bind CD1B tetramer loaded with GMM (Figure 3A, 3B and Supplementary Figure 1). The clonotype diversity observed after in vitro expansion was qualitatively similar to that observed in ex vivo samples (Figure 3A and Supplementary Figure 1). For example, there was marked enrichment of TRAJ26 in the ex vivo tetramer-positive and expanded and resorted populations compared to tetramer-negative cells (Figure 3A). A similar pattern was observed within the TCR-β chain sequences, in which tetramer-positive cells showed a clear bias towards the use of certain variable and joining segments as compared to those observed in the tetramer-negative cells (Figure 3B and Supplementary Figure 1). We then compiled lists of the most highly enriched V and J gene segments from TCR-α (TRAV1, TRAJ9) and TCR-β (TRBV6, TRBJ2) by comparing expanded and resorted tetramer-positive cells to tetramer-negative cells and analyzing the length of the CDR3 region. We found that CDR3 length was severely restricted in these cell populations (Figure 3C and Supplementary Figure 2). Collectively, these data provide further evidence of substantial restriction in the TCR-α and TCR-β chains that bind CD1B-GMM tetramers.

**Figure 3.**
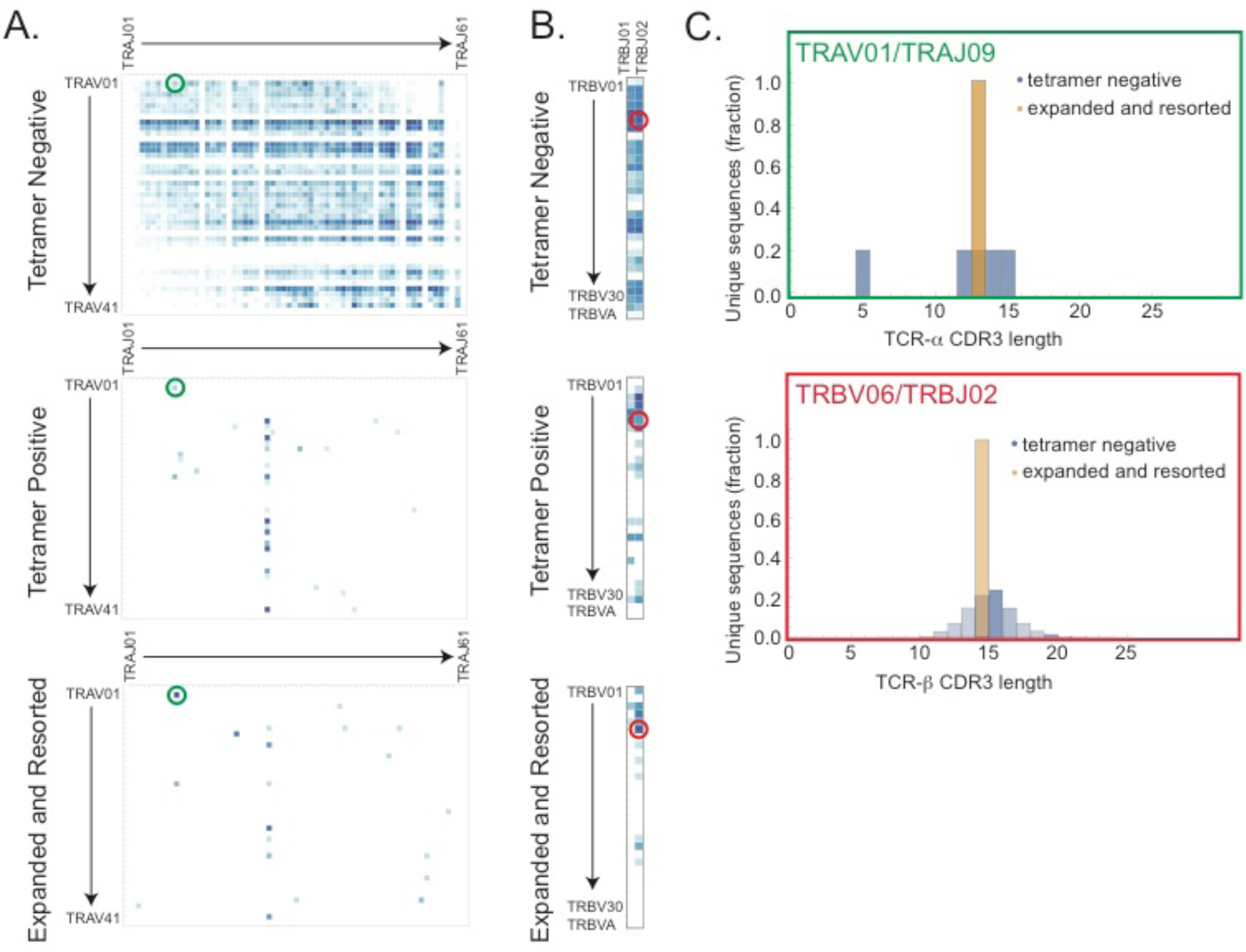
GMM-specific T-cell receptor diversity. Heatmaps depicting the diversity of (A) TRAV/TRAJ or (B) TRBV/TRBJ recombination events found among tetramer-negative, tetramer-positive, or expanded and resorted T cells obtained from one representative blood donor with latent tuberculosis infection. Blue color scale indicates the relative abundance of each V/J combination. The green circle represents TRAV1-2 and TRAJ9 rearrangement. The red circle represents the TRBV6 and TRBJ2 rearrangement (C) Histograms show the distribution of CDR3 length among the most highly enriched V and J gene segment combinations for TCR-α (TRAV1/TRAJ9) and TCR-β (TRBV6/TRBJ2).

### Unbiased definition of a shared GMM-specific TCR motif

To determine whether there is a GMM-specific T-cell repertoire that is shared among unrelated donors, we standardized the analysis described above by constructing a simple test to identify TCR motifs. We defined motifs as sequences with common V and J family usage and CDR3 length that were significantly enriched in the expanded and re-sorted samples, as compared to their corresponding tetramer-negative samples. This unbiased approach resulted in one TCR-α motif and one TCR-β motif that were significantly enriched independently in each of the four donors. The TCR-α motif contained a 13-amino acid CDR3 sequence that is consistent with the recently published TRAV01/TRAJ09 rearrangement present in GEM T cells (Figure 4A and Supplementary Table 1) (15). The TCR-β motif we discovered in this manner included a 14-amino acid CDR3 sequence that results from a TRBV06/TRBJ02 rearrangement (Figure 4B and Supplementary Table 1). Furthermore, several lines of evidence support co-expression of these motifs as a heterodimeric TCR in unrelated individuals. First, the dominant TCR-α and TCR-β sequences for one of our T cell lines analyzed in this study matches these motifs, suggesting they are co-expressed by a dominant T-cell clone (data not shown). Second, when we sorted GMM-CD1B tetramer avid cells from a fifth unrelated South African donor and performed limiting dilution to establish T-cell clones, we identified two T-cell clones whose TCRs match the TCR-α and TCR-β motifs described above (Supplementary Table 2). Finally, Van Rhijn et al. reported a GMM-specific T-cell clone derived from an M.tb-infected subject whose TCR also matches these TCR-α and TCR-β motifs (15). Taken together, these data suggest that specific TCR-α and TCR-β chains that conform to the motif described above are co-expressed as a heterodimer, are specific for the mycobacterial antigen GMM, and are shared among multiple unrelated donors.

**Figure 4.**
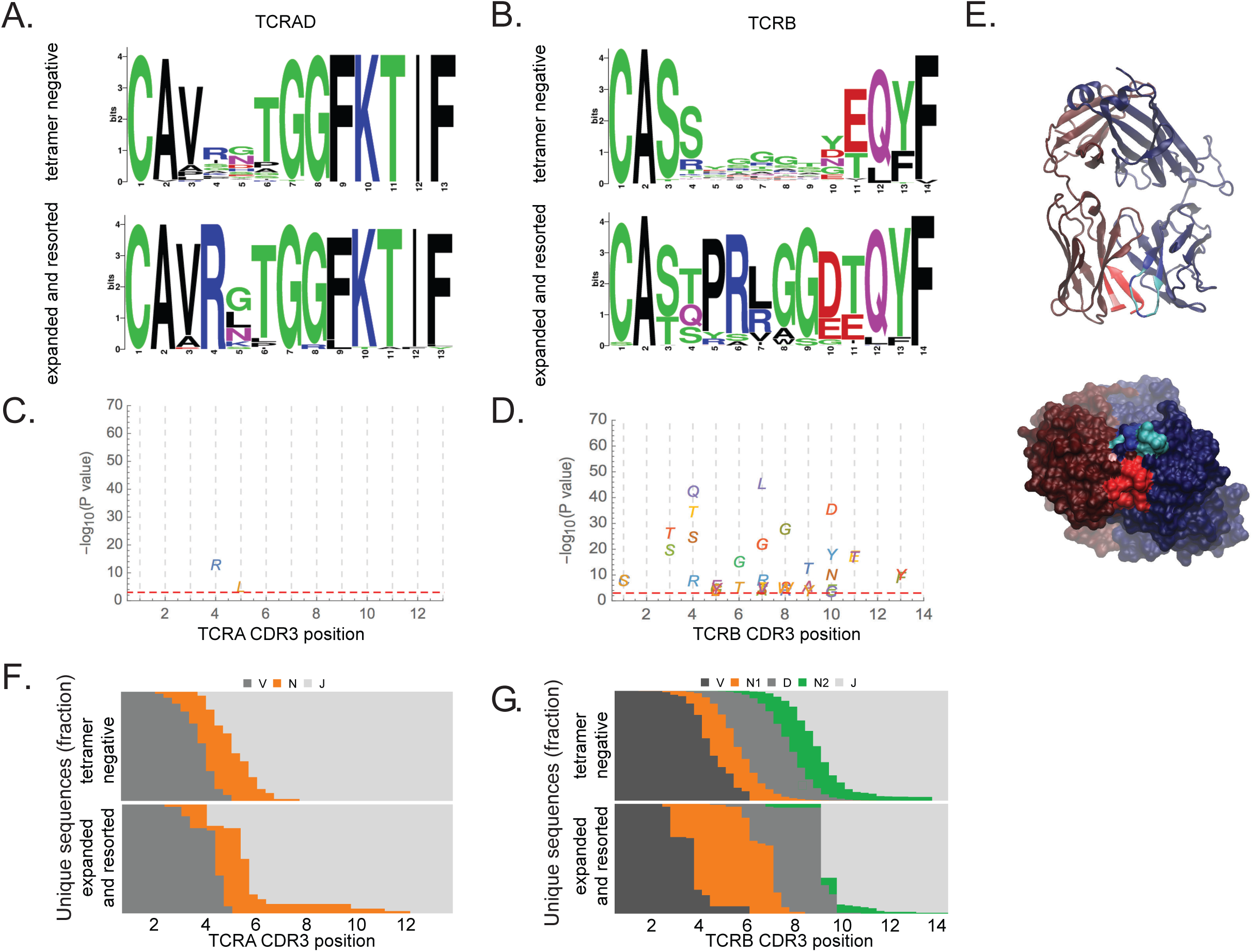
Conservation among sequences matching the CD1B-GMM specific TCR motifs. Logo plots indicating the positional CDR3 amino acid usage among sequences within (A) TRAV01 and TRAJ09 rearrangement for TCR-α and (B) TRBV06 and TRBJ02 rearrangement for TCR-β among expanded and resorted samples as well as tetramer-negative samples. At each position along the CDR3 region, the statistical significance of enrichment of each residue in the expanded and resorted samples as compared to tetramer-negative samples is indicated for (C) TCR-α and (D) TCR-β. The selected significance threshold of p-value = 10^−3^ is indicated with a red dashed line. (E) The crystal structure of a published CD1B-GMM specific TCR whose sequence matches both the TCR-α and TCR-β motifs detected in this study. The TCR-α CDR3 is indicated in red, with position of significantly enriched residues indicated in pink. The TCR-β CDR3 is indicated in blue, with position of significantly enriched residues indicated in cyan. A portion of the TCR-α CDR3 was not resolved in the crystal structure, resulting in an apparent gap. At each position along the (F) TCR-α or (G) TCR-β CDR3 region, the proportion of sequences annotated as originating in a germline V, J, or D gene, as well as non-templated (N) bases at that position is depicted.

### Predicted binding interactions and ontogeny of the GMM-specific TCR motifs

We next addressed the question of how, given the stochastic nature of T-cell development, these motifs might arise independently in unrelated donors. To accomplish this, we first assessed the enrichment of particular amino acid residues at each position along the CDR3 of expanded and re-sorted samples. In this analysis, we used tetramer-negative samples as a natural control and looked for differences in the CDR3 sequence of the motifs described above. In the TCR-α motif, we observed that the arginine at position 4 and leucine at position 5 were enriched, suggesting that these residues are particularly important for mediating binding to the GMM-CD1B tetramer (Figure 4C and Supplementary Tables 3 and 4). This inference is supported by TCR reconstitution experiments, in which a TCR containing a TCR-α chain that lacks arginine at position failed to respond functionally to GMM (15). We also identified several enriched residues between positions 4 and 13 of the TCR-β motif (Figure 4D and Supplementary Table 4). Figure 4E shows the location of these enriched residue positions identified through sequence analysis mapped to the crystal structure of a GMM-specific TCR whose TCR-α and TCR-β chains match our motifs (15). Interestingly, both chains of the heterodimer appear to contribute equally to binding the CD1B-GMM tetramer, which is supported by the recently published co-crystal of a GEM TCR with CD1B-GMM (30). These data highlight how immunosequencing can be used to infer structural requirements for antigen binding by TCRs.

We next explored the distribution of nucleotide additions and deletions leading to junctional diversity in the TCR-α motif described above. We observed a lower level of 3’ deletion in the V segment of TCR-α sequences from expanded and resorted T-cells than in the those from tetramer-negative cells (Figure 4F). We note that the germline definition for TRAV01 ends with CAVR (Figure 4F), which includes the conserved arginine at position 4 required for binding CD1B-GMM, suggesting that this CDR3 is germline-encoded, as previously proposed (15). In contrast, the germline sequence of TRAJ09 begins with GNTGGFKTIF, implying that position 5 of the selected motif results from selection of non-templated insertions that result in the encoding of a leucine residue in CD1B-GMM specific TCRs. Alternatively, it is possible that modulation of the residue distribution at position 5 could be a consequence of the restriction observed at position 4, since non-templated insertion has been shown to be context dependent (31). When we performed the same analysis for TCR-β, we observed an increased level of 3’ V deletion in CD1B-GMM specific TCRs as compared to those observed in TCRs from tetramer-negative cells, suggesting that enriched residues at positions 4, 5, and 6 are likely non-templated. On the other hand, the enriched residues at positions 10 to 13 are located entirely within the germline-encoded TRBJ02 gene segment (Figure 4G). These data show that a GMM-specific shared TCR could arise through a combination of germline rearrangements and high-probability events occurring at junctions.

### Ex-vivo analysis of GMM-specific TCR motifs in healthy donors

To extend the results described above, which were obtained from *in-vitro* expanded and resorted T-cell lines, we next examined sequences obtained from tetramer-positive cells sorted directly *ex vivo* (Figure 2A). We pooled sequences from all four subjects studied initially and considered sequences concordant with the previously defined TCR-α and TCR-β motifs in terms of V and J gene family and CDR3 length as matches (Figure 4A and 4B). We observed a significant enrichment of both the TCR-α and TCR-β motifs among tetramer-positive cells as compared to tetramer-negative cells (Table 1). These data support the utility of the TCR motifs identified in this study as markers of antigen-specific T-cell responses *ex vivo*. We next examined the prevalence of the TCR-β motif in a previously published dataset that included TCR-β sequences obtained from the PBMC of 587 bone marrow donors at low risk for *Mycobacterium tuberculosis* infection (32). We found that the motif was present at a frequency very similar to that observed in tetramer-negative T cells in the pooled data (Table 1). The relative absence of the GMM-specific TCR-β motif in T cells from healthy donors suggests that it could be used as a specific marker of mycobacterial infection in population-based studies.

**Table 1.** **Frequency of CD1B-GMM specific TCR motifs among sorted cells and PBMC from bone marrow donors.** We pooled unique sequences from all individuals for each sample type and each locus and counted the number matching (or mismatching) the defined CD1B-GMM specific TCR motifs. Tetramer-positive and expanded and resorted samples exhibit significant enrichment of motifs in both TCR-α and TCR-β as compared to tetramer-negative samples. PBMC from 587 healthy bone marrow donors have a similar number of TCR-β motif matches as tetramer-negative samples. P-values were computed with Fisher’s Exact test.

|  | Sequence Source | Motif Match | Motif Mismatch | Match/Mismatch | Fold Enrichment | p-value |
| --- | --- | --- | --- | --- | --- | --- |
| TCR-α | Tetramer-negative | 55 | 174409 | 0.0003152 | | |
| | Tetramer-positive | 52 | 573 | 0.0832 | 263.92 | $6.21 \times 10^{-98}$ |
| | Expanded & resorted | 46 | 123 | 0.2721 | 863.40 | $3.34 \times 10^{-113}$ |
| TCR-β | Tetramer-negative | 7613 | 462440 | 0.01619 | | |
| | Tetramer-positive | 126 | 566 | 0.1821 | 11.24 | $9.88 \times 10^{-89}$ |
| | Expanded & resorted | 119 | 255 | 0.3182 | 19.65 | $5.50 \times 10^{-115}$ |
|  | Bone marrow donors | 2172898 | 130501755 | 0.0164 | 1.01 | 0.17 |

### GMM-specific TCR motifs during tuberculosis

Finally, we investigated the association between GMM-specific TCR motifs and mycobacterial infection by analyzing a cohort of South Africans with known M.tb infection status. This cohort included three clinical groups: adolescents without latent tuberculosis infection (IGRA-negative), adolescents with latent tuberculosis (IGRA-positive), and adults with a new diagnosis of active tuberculosis disease (Active TB, n=10). We comprehensively profiled the TCR-α and TCR-β chains from an average of approximately 100,000 T cells per donor by immunosequencing cryopreserved PBMCs, and we pooled the data into each of the three clinical groups. Next, we calculated and compared the ‘motif burden’ of the TCR-α and TCR-β motifs described above, defined as the fraction of unique sequences that matched the V and J gene family, and CDR3 length of these motifs (Table 2). For the TCR-α motif, we noted an increased motif burden in IGRA-positive subjects (p=0.0014) and active TB patients (p=8.3 × 10^7^) compared to IGRA-negative control subjects. For the TCR-β motif, we also observed an increased motif burden in IGRA-positive subjects (p=4.9 × 10^−5^), and active TB patients (p=8.3 × 10^−7^) when compared to IGRA-negative subjects (Table 2). When we calculated subject-specific motif burdens, we confirmed enrichment in the TCR-α when comparing active TB patients to IGRA-negative subjects (p=0.004), although in this case we did not see an enrichment in the TCR-β motif burden (Figure 5A and Supplementary Figure 3). Notably, we did not see enrichment of canonical MAIT cell or iNKT cell TCR-α rearrangements in active TB, highlighting the specificity of the association between a GMM-specific TCR-α motif and tuberculosis (Supplementary Figure 3).

**Figure 5.**
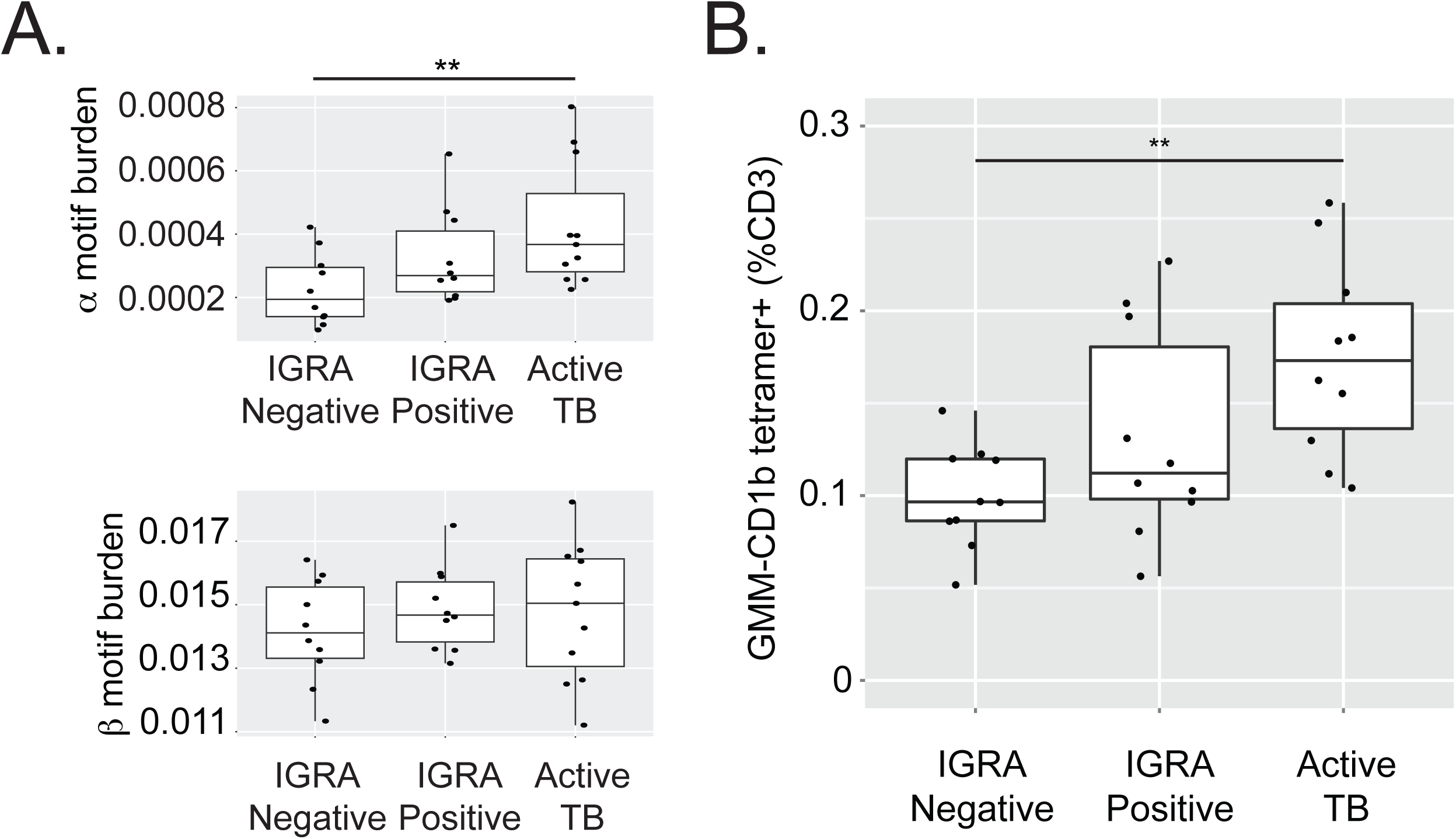
GMM-specific TCR motifs and T cells are increased in frequency during tuberculosis infection and disease. For each subject, we calculated the fraction of TCRs matching the motifs defined in this study that were present in the peripheral blood of IGRA-negative, IGRA-positive, and active TB patients. (A) Within TCR-α, there is an increased motif burden in active TB subjects compared to IGRA-negative subjects (Kruskal-Wallace p=0.03, Dunn post-test p=0.0038). There was not a statistically significant difference in subject-specific TCR-β motif burdens among the three groups (Kruskal-Wallace p=0.67). (B) The frequency of GMM-CD1B tetramer positive T cells by flow cytometry is increased in patients with active TB as compared to IGRA-negative subjects (Kruskal-Wallace p=0.01, Dunn post-test p=0.0013).

**Table 2.** **Frequency of CD1B-GMM specific TCR motifs among M.tb-uninfected and M.tb-infected South African subjects.** We performed immunosequencing for each locus from IGRA-negative (n=10), IGRA-positive (n=10), and active TB (n=10) subjects, and then pooled the sequences for each locus from all subjects with a specific infection status. We tabulated the number of sequences in the pool matching the defined CD1B-GMM specific TCR motif, as well as the complementary number of sequences not matching the motif. Both IGRA-positive subjects and patients with active TB exhibit significant enrichment of both TCR-α and TCR-β motifs as compared to IGRA-negative subjects. P-values were computed using Fisher’s Exact test on the raw counts of matches and mismatches. Because sampling depths varied widely among subjects, this pooling approach complements the subject-wise analysis included in Fig. 5b, which eliminated sampling depth information by computing fractional motif burdens

|  | Sequence Source | Motif Match | Motif Mismatch | Match/Mismatch | Fold Enrichment | p-value |
| --- | --- | --- | --- | --- | --- | --- |
| TCR-α | IGRA-negative | 180 | 817804 | 0.0002201 | | |
| | IGRA-positive | 228 | 765928 | 0.0002976 | 1.35 | $1.40 \times 10^{-3}$ |
| | Active TB | 224 | 628088 | 0.0003565 | 1.62 | $8.29 \times 10^{-7}$ |
| TCR-β | IGRA-negative | 7565 | 532381 | 0.01401 | | |
| | IGRA-positive | 8233 | 544217 | 0.01490 | 1.06 | $4.86 \times 10^{-5}$ |
| | Active TB | 8495 | 563097 | 0.01486 | 1.06 | $8.75 \times 10^{-5}$ |

We further validated these results using flow cytometry to quantify GMM-specific T cells ex vivo (Supplementary Figure 4). We found that the frequency of CD3+ T cells that stained with CD1B-GMM tetramer was higher in patients with active TB as compared to IGRA-negative control subjects (p=0.001, Figure 5B). Finally, in one patient with active tuberculosis, we analyzed blood samples at diagnosis, during treatment, and at the completion of therapy, and we observed a decrease in the frequency of sequences that matched both the TCR-α and TCR-β motifs over time (Supplementary Figure 3). Together, these data support clonal expansion and contraction of GMM-specific T cells during mycobacterial infection and clearance, respectively, and demonstrate the association between GMM-specific TCR motifs and M.tb infection and disease.

## DISCUSSION

The phenomenon of MHC restriction has been a cardinal feature of T-cell immunology for more than forty years. In practical terms, this means that the ability to activate T cells depends not only on the foreign antigens they recognize, but also on the MHC molecules that bind the antigen. By contrast, we show here that CD1B is nearly invariant among humans, and we report T-cell receptor motifs that are specific for a mycobacterial lipid antigen presented by CD1B, and consistently shared among multiple unrelated donors. We also show that T cells bearing these motifs are more frequent in South Africans with latent M.tb infection or tuberculosis disease than in M.tb-uninfected subjects. These data support the existence of a lipid antigen-specific shared T-cell repertoire that is independent of genetic background and could be leveraged to develop molecular diagnostics based on TCR sequence motifs.

There are numerous examples of ‘public’ TCR sequences that are shared across unrelated donors. Most of these have been reported as reactive to viral peptide antigens (reviewed in (33) and (34)). However, these reports also make it clear that most T-cell responses are ‘private’ to a single individual, and that public T-cell responses are the exception rather than the rule (7). Importantly, since most of these TCRs likely bind peptide-based epitopes, they are by definition MHC-restricted. On the other hand, some T cells that share an invariant TCR-α chain, such as iNKT and MAIT cells, are widely prevalent but not clearly antigen-specific or related to pathogen exposure. In this study, we show that for CD1B presentation of GMM, a shared antigen-specific TCR response is seen reliably in more than 30 unrelated individuals. Our data support a model that requires a conserved, but not necessarily invariant, rearrangement in both α and β chains of the TCR heterodimer to recognize a foreign lipid antigen. Additionally, by comparing the TCR sequences of GMM tetramer-positive and tetramer-negative T cells, we provide a structural rationale for this notion that is confirmed by the recent publication of GEM T cells and associated crystal structures (15, 30). Our data support the emerging paradigm that TCR recognition of foreign lipids follows a ‘parallel docking mode’ that is more reminiscent of peptide-MHC than of CD1d-αGalCer TCR interactions, which is relatively biased toward the TCR-α chain (35).

Our unbiased approach to motif discovery is thus validated in the setting of GMM, suggesting it can be applied more broadly to lipid antigens outside of infectious diseases. For example, T cells have been described that recognize methyl-lyso phosphatidic acid when bound to CD1C expressed by acute myeloid and B cell leukemias (36). T cells recognizing lysophospholipids bound to CD1A contribute to pathogenic inflammation in atopic dermatitis and psoriasis (37, 38). Human CD1A and CD1C tetramers as well as synthetic lipid antigens are available as validated reagents, so that an approach similar to the one described here could potentially be used to define allergic or cancer-specific shared TCRs and to test their clinical associations.

We detected GMM-specific TCR motifs in subjects without M.tb infection, suggesting either a high frequency of these motifs in the naïve T cell repertoire as a consequence of near-germline rearrangements, T-cell priming through mechanisms other than infection with *M. tuberculosis*, or both. Because GMM is produced by mycobacteria other than M.tb, this phenotype could result from exposure to non-tuberculous mycobacteria in the environment, or from vaccination with the *M. bovis* derived-BCG vaccine at birth, which is standard practice in South Africa. Consistent with this possibility, we found that the TCR-β motif was not enriched in a cohort of U.S. bone marrow donors who are at low risk for M.tb exposure and did not receive BCG. We can apply our unbiased approach to identify shared TCRs for lipids that are preferentially expressed by virulent M.tb. Unlike GMM, sulfoglycolipids are not expressed by BCG or most environmental bacteria, and T-cell responses to sulfoglycolipids have been shown to be greater in M.tb-infected subjects when compared to BCG-vaccinated subjects (39). The discovery of shared TCRs covering a variety of mycobacterial lipids could then be tested independently or in combination for utility as a molecular marker for tuberculosis.

We used both immunosequencing and flow cytometry to demonstrate that GMM-specific T cells in peripheral blood are expanded during active tuberculosis. The increased abundance of the GMM-specific TCR motifs we identified among active TB patients suggests expansion of high-avidity clones in the presence of antigen. Moreover, we show a decrease in GMM-specific TCRs during treatment for active tuberculosis. These dynamic changes are consistent with the development of immunologic memory to lipid antigens. By contrast, we show that the frequency of TCRs expressed by innate-like T cells, such as iNKT and MAIT cells, are not associated with active tuberculosis. In this respect, GMM-specific T cells may more closely resemble conventional MHC-restricted T cells rather than innate-like T cells. Future studies could address whether these TCRs specific for mycobacterial lipids could be used as molecular markers of response to drug treatment or immunogenicity of whole cell vaccines, such as BCG.

The advantage of maintaining a highly polymorphic pool of MHC genes in an outbred population have been extensively explored (40). In contrast, the advantages of maintaining a nearly invariant CD1B gene are less obvious. While MHC finds its origins in jawed vertebrates, CD1 genes have only been reported in mammals and birds, suggesting CD1 evolved from MHC genes to perform a non-redundant function. A number of pathogens have evolved mechanisms to avoid detection by the adaptive immune system, by varying the sequence of peptides available to bind host MHC molecules (41). Thus, CD1 may have evolved to facilitate T-cell detection of non-peptide antigens that are less likely to undergo mutational escape because of their importance in maintaining cell wall integrity. Supporting this notion is the success of mycolic acid synthesis inhibitors, such as isoniazid, for the treatment of tuberculosis. Isoniazid is predicted to also inhibit the production of GMM (42). Whether the findings we report here can be generalized to other lipids or non-peptide antigens remains to be determined, as do the true evolutionary origins of non-MHC systems of antigen presentation.

In summary, we provide evidence of purifying selection in human CD1B and identify a set of conserved TCRs specific for CD1B and a mycobacterial lipid that is enriched during tuberculosis infection and disease. These data demonstrate a general framework for the rapid and unbiased identification of T cells specific for CD1-presented antigens as well as their associations with disease. These T cells are part of a potentially much larger shared T-cell repertoire that might have a number of potential clinical applications. One possibility is molecular diagnosis of M.tb or any other disease characterized by lipid antigen-specific T-cells, such as atopic dermatitis, psoriasis, or leukemia. The other possibility is adoptive cell therapies that do not depend upon the genetic background of the patient. This is in contrast to current clinical trials of TCR-modified T cells, for example, that are typically limited to the dominant MHC haplotypes present in a population at risk for cancer (43). The realization of such novel cell therapies will require the identification of disease-specific lipid antigens and shared TCRs, but much of the technology for translating this information into a clinical therapy is already available.

## MATERIALS AND METHODS

### Generation of GMM-loaded CD1B tetramers

Soluble biotinylated CD1B monomers were provided by the National Institutes of Health Tetramer Core Facility (Emory University, Atlanta, GA). Glucose monomycolate (GMM)-loaded tetramers were generated as previously described (29). In brief, C32-GMM was dried down in a glass tube using a nitrogen evaporator and sonicated into 0.25∼ CHAPS/sodium citrate at pH 4 (preparation of CHAPS in sodium citrate; CHAPS Hydrate, Sigma; Sodium Citrate Dihdrate, Fisher) for two minutes at 37°C. The lipid solution was transferred to a microfuge tube, and 9 μL of CD1B monomer was added. The CD1B-GMM preparation was then incubated in a 37°C water bath for 2 hours with vortexing every 30 minutes. At the end of the incubation, the solution was neutralized to pH 7.4 with 6 μL of 1M Tris pH 9. Finally, 10 μL of Streptavidin conjugated to allophycocyanin or phycoerythretin (Life Technologies) was added in ten aliquots of 1 μL every 10 minutes to facilitate tetramerization. The final product was filtered through a SpinX column (Sigma) to remove aggregates and stored at 4°C until use.

### Antigens and Media

C32-GMM purified from *Rhodococcus equi* was generously provided by the laboratory of D. Branch Moody (44). Our base T cell media consists of RPMI 1640 (Gibco) supplemented with 100 U/mL Penicillin and 100 μg/mL Streptomycin (Gibco), 55 μM 2-mercaptoethanol (Gibco),0.3X Essential Amino Acids (Gibco), 60μM Non-essential Amino Acids (Gibco), 11mM HEPES (Gibco), and 800μM L-Glutamine (Gibco). We additionally supplemented this media with either fetal calf serum (HyClone) or human serum that was derived from healthy donors.

### Generation of T cell lines using GMM-loaded CD1B tetramers

Peripheral blood mononuclear cells (PBMC) were isolated from South African adults with latent tuberculosis infection as defined by a positive QuantiFERON-TB Gold blood test. PBMC were depleted of CD14-expressing monocytes for a separate study and cryopreserved. At the time of this study, PBMC were thawed and washed in warm RPMI 1640 (Gibco) supplemented with 10∼ fetal calf serum (Hyclone) and 10 μL/mL Benzonase (Millipore) and enumerated using Trypan blue exclusion. PBMC were then plated in a 24-well plate at a density of three million cells per well in T cell media and allowed to rest overnight at 37°C in humidified incubators supplemented with 5∼ CO_2_. The following day, PBMC were washed and blocked with 50∼ human serum (Valley Biomedical) in PBS supplemented with 0.2∼ BSA (Sigma) (FACS buffer) for 10 min at 4°C. The samples were washed twice with PBS and stained with Aqua Live/Dead stain (Life Technologies) according to the manufacturer’s instructions. Following two additional PBS washes, cells were resuspended in 50 μL FACS buffer and 1 μL of either unloaded CD1B tetramer or GMM-loaded CD1B tetramer and incubated at room temperature for 40 minutes in the dark. Finally, cells were stained with anti-CD3 ECD (Beckman Coulter) as well as surface markers for activation and memory phenotype, washed twice in T cell media, and screened through a cell strainer tube (Falcon) prior to sorting. Tetramer-positive T cells were sorted at the UW Department of Immunology Flow Cytometry Core using a FACS Aria II (BD) cell sorter equipped with 407nm, 488nm, and 641nm lasers.

Sorted T cells were washed and resuspended in T cell media supplemented with 10∼ human serum. T cells were then divided among eight wells of a 96-well U-bottom tissue culture plate into which irradiated PBMC (150,000 cells per well) were added as feeder cells along with phytohaemagglutinin (Remel) at a final concentration of 1.6 μg/ml. After two days in culture at 37°C, 5∼ CO_2_, 10 μL natural IL-2 (Hemagen) was added to each well. Half the media was replaced every two days with T cell media supplemented with 10∼ human serum and natural IL-2 When the cell clusters were large and round (approximately after eight days of growth), they were pooled into a 24-well plate. After 10 days in culture, cell lines were screened by tetramer staining or functional response to GMM. We then further expanded T cell lines by modifying a previously published rapid expansion protocol (45). Briefly, 200,000 T cells were mixed with 5 million irradiated EBV-transformed B cells and 25 million irradiated PBMC as feeder cells in T cell media. Anti-CD3 (clone OKT3) was added a final concentration of 30 ng/mL, and the mixture was incubated overnight at 37°C, 5∼ CO_2_. The following day, recombinant IL-2 (rIL-2) (UWMC Clinical Pharmacy) was added to a final concentration of 50 U/mL. On day 4, the cells were washed twice in T cell media to remove OKT3, and fresh media supplemented with rIL-2 at 50 U/mL was added. Half the media was replaced every three days or split into new T25 tissue culture flasks (Costar) as determined by cell confluency. After 13 days in culture, the lines were screened by tetramer staining and then cryopreserved on day 14.

### Clinical Cohorts

As recently published, 6363 adolescents were enrolled into a study that aimed to determine the incidence and prevalence of tuberculosis infection and disease (46). Twelve to 18 year-old adolescents were enrolled at eleven high schools in the Worcester region of the Western Cape of South Africa. Subjects were screened for the presence of latent tuberculosis by a tuberculin skin test and IFN-γ release assay (IGRA) QuantiFERON-TB GOLD In-Tube (Cellestis Inc.) at study entry. Peripheral blood mononuclear cells (PBMC) were isolated from freshly collected heparinized blood via density centrifugation and cryopreserved. For this work, a subset of samples from 10 M.tb-infected and 10 M.tb-uninfected adolescents were selected based on matching for age and sex and availability of cryopreserved specimens after completion of the primary objectives of the parent study. We also accessed a recently published cohort of South African adults with a new diagnosis of active tuberculosis (47). Only participants were included when ≥ 18 years of age and seronegative for HIV were included. All included patients had either positive sputum smear microscopy and/or positive culture for *M.tb*. Blood was obtained and PBMC archived prior to or within 7 days of starting standard course anti-TB treatment, which was provided according to South African national health guidelines. For this work, a convenience sample from 10 adults with active tuberculosis was selected based on availability of cryopreserved specimens after completion of the primary study. For T-cell cloning studies, we used cryopreserved PBMC from a cohort of healthy South African adults (Supplemental Table 5).

Human peripheral blood samples were also obtained from a cohort of healthy bone marrow donors from the Fred Hutchinson Cancer Research Center Research Cell Bank biorepository, under a protocol approved and supervised by the Fred Hutchinson Cancer Research Center Institutional Review Board, following written informed consent. This cohort has been previously described (32) (Emerson et al., manuscript submitted).

### Flow Cytometry

PBMC from South African adolescents and adults were thawed and rested overnight as described above. The following day, one million PBMC per subject were counted and plated into each of two wells of a 96-well U-bottom plate. The cells were spun down and blocked in human serum and FACS buffer at 1:1 for ten minutes at 4°C. Following a wash with FACS buffer, the cells were resuspended in 50 μL FACS buffer, and 1 μL of either unloaded CD1B tetramer conjugated to PE or 1 μL GMM-loaded CD1B tetramer conjugated to APC. The samples were incubated at room temperature for 40 minutes and then washed twice with PBS. Aqua Live/Dead stain was then prepared at a 1:100 dilution in PBS and added to each sample. All samples were incubated for 15 min at room temperature in the dark. The samples were then washed twice and resuspended in 50 μL of staining cocktail containing anti-CD3 ECD in FACS buffer for 30 minutes at 4°C in the dark. Cells were then washed twice in FACS buffer and centrifuged at 1800 rpm for 3 min. All samples were then resuspended in 200 μL of 1∼ paraformaldehyde prior to acquisition on LSRII (BD Biosciences) equipped with 488nm (100mW), 532nm (150mW), 628nm (200mW), 405nm (100mW), and 355nm (20mW) lasers.

### Functional Assays

We used IFN-γ ELISPOT to examine antigen-specificity of T cell lines. Briefly, K562 cells stably transfected with empty vector (K562-EV) or CD1B (K562-CD1B)(48) were maintained in RPMI 1640 (GIBCO) supplemented with 10∼ fetal calf serum and G418 (Sigma) at a concentration of 200 μg/ml and periodically assessed for CD1B expression by flow cytometry. Multiscreen-IP filter plates (Millipore) were coated with 1-D1K antibody (Mabtech) overnight at 4°C. Lipids were evaporated to dryness from chloroform-based solvents under a sterile nitrogen stream and then sonicated into media. This lipid suspension was added to co-cultures of 50,000 mock transfected or CD1B-transfected K562 antigen-presenting cells and 2000 T cells with a final concentration of 1 μg/ml for MA and GMM (8, 49). The co-cultures were incubated for 16 hours at 37°C. The following day, the cells were lysed with water and then incubated with 7-B6-1 biotin conjugate (Mabtech) for two hours at room temperature. The plate was washed with PBS (Life Technologies) and then incubated with ExtraAvidin-Alkaline Phosphatase (Sigma) for 1 hour at room temperature. Lastly, the wells were washed and then developed using BCIP/NBT substrate (Sigma) for 5 min at room temperature in the dark. The wells were imaged and the IFN-γ spots were counted using an ImmunoSpot S6 Core Analyzer (Cellular Technology Limited).

To observe intracellular cytokine staining following GMM antigen stimulation, 3.3 million K562 cells/ml were incubated with 5μg/mL GMM at a final volume of 100 μ! for 18 hours at 37°C, 5∼ CO_2_ to facilitate lipid loading. The following day, T cell lines were plated at 1 million cells/mL and 80μL of pre-loaded K562 cells was added to the T cells. The cell mixture was allowed to incubate for six hours in the presence of anti-CD28/49d antibodies (BD Biosciences), Brefeldin A at a final concentration of 10 μg/ml (Sigma), and GolgiStop containing Monensin (BD), and anti-CD107a (BD) after which EDTA, at a final concentration of 2mM, was added to disaggregate cells. Plates were stored at 4°C until the following day when they were stained and acquired by FACS. We used a previously published optimized and validated 12-color panel (50, 51). Briefly, cells were first stained with Avid Live/Dead (Life Technologies) viability dye and anti-CD14. After washing, the cells were permeabilized with FACS Perm II (BD) and stained for the remaining markers (CD3, CD4, CD8, CD154, IFN-γ, TNF-α, IL-2, IL-4, IL-17a, and CD40L). Fully stained cells were washed and resuspended in 1∼ paraformaldehyde. Data were acquired on an LSR II (BD) equipped with a high-throughput sampler and configured with 405nm (50mW), 488nm (20mW), 561nm (50mW), and 641nm (150mW) lasers.

### Ethics

The study was approved by the IRB of the University of Washington and the University of Cape Town. Written informed consent was obtained from all adult participants as well as from the parents and/or legal guardians of the adolescents who participated. In addition, written informed assent was obtained from the adolescents.

### Comparative Genomics

Data for CD1B (Chr1:158,327,951-158,331,531) and HLA-A (Chr6:29,941,260-29,945,884) were extracted from the 1000 Genomes Project (Phase 3; aligned to GRCh38), and measures of divergence calculated using VCFtools. Nucleotide diversity (π) (http://www.pnas.org/content/76/10/5269.abstract) was estimated with sliding windows of 50 base pair with two base pair steps, and Tajima’s D (https://www.ncbi.nlm.nih.gov/pmc/articles/PMC1203831/) was estimated over static 50 base pair windows. Coding SNP variation with >1∼ minor allele frequency was extracted, and manually mapped onto the corresponding protein structures (CD1B: PDB #1UQS, HLA-A: PDB #3MRG) according to variant type (synonymous or nonsynonymous) (24, 27).

### Immunosequencing

High-throughput sequencing of TCRs was performed using the ImmunoSEQ assay (Adaptive Biotechnologies) with TCR-β and/or TCR-α/δ assays for each sample using a multiplex PCR approach following by Illumina high-throughput sequencing (52).

### Putative CD1B-GMM specific TCR motifs defined by V/J gene usage and CDR3 length

Sequence motifs for CD1B-GMM specific TCR-α and TCR-β were defined for each subject by assessing enrichment of specific V and J gene family usage and CDR3 lengths within the expanded and resorted sample as compared to the respective tetramer negative sample. For each possible combination of V family, J family, and CDR3 length, we enumerated unique sequences matching this combination, as well as the mismatching sequences for each sample. From these four counts (expanded/resorted, tetramer-negative, matching, and mismatching), we generated a 2×2 contingency table and used Fisher’s Exact test to assign a p-value for the significance of the enrichment of matches in the expanded and resorted sample. Such a test was performed for all combinations for each subject and for both TCR-α and TCR-β data, and we selected those with p-values below a pre-specified significance threshold (p < 0.001). This resulted in a set of CD1B-GMM specific TCR motifs for each subject. To define the ‘public’ CD1B-GMM specific TCR motifs within each locus, we identified motifs that were significantly enriched as described above among all subjects.

### Sequence conservation among CD1B-GMM specific TCRs matching putative motifs

We determined the sequence conservation among putative motif matches of expanded and resorted TCRs (i.e. considered truly CD1B-GMM specific),s using putative motif matches among sequences derived from tetramer-negative cells as a natural control. For each possible amino acid at a given position in the CDR3, sequences matching the putative motif that contained this amino acid at the given position were enumerated for expanded and resorted data and tetramer-negative data, as were the sequences not matching that specific amino acid at the given position. From these four counts, we generated a 2x2 contingency table and used Fisher’s Exact test to assign a p-value for significance of enrichment of this amino acid at the given position among the expanded and resorted putative motif matches. Such a test was performed for amino acids and CDR3 positions and for both TCR-α and TCR-β data, and we selected those with p-values below a pre-specified significance threshold (p-value < 0.001). Using the annotations of V, N, and J region boundaries (for TCR-α sequences) and V, N1, D, N2, and J region boundaries (for TCR-β sequences), we were also able to assess differences in recombination structure among CD1B-GMM specific TCRs with respect to nonspecific TCRs that also matched the putative motifs (53). By integrating these observations with the positions and identities of conserved residues, we were able to assess the conservation of germline-encoded sequence versus specific non-templated insertions.

### CD1B-GMM specific TCR motif burden

For population-level comparisons between T-cell populations or between patient groups, we defined the ‘motif burden’ of a sample simply as the fraction of unique TCRs from that sample that matched the CD1B-GMM specific TCR motif defined for the corresponding TCR-α or TCR-β. We first assessed significance by pooling all the sequences within each clinical group and counting the number that matched and the number that mismatched each motif using Fisher’s exact test on the resulting contingency table. Pooling sequences from multiple donors is sensitive to one donor with an excess of sequences biasing the result. To address this, we examined the sampling depth across donors in in each clinical group, and found that healthy subjects were not systematically undersampled (data not shown). We also assessed differences between clinical groups using subject-specific fractional motif burdens using the non-parametric Kruskall-Wallace test with Dunn post test (a generalization of the Mann-Whitney U test to more than two groups).

## AUTHOR CONTRIBUTIONS

C.S. and R.O.E. designed the study. W.S.D. and K.K.Q performed the experiments and analyzed the data. D.W. and W.S. contributed human genetic analyses. C.L.D. and T.J.S. established the clinical cohorts. S.C.D facilitated flow cytometry studies. A.S. and H.S.R. facilitated immunosequencing. W.S.D, M.V., R.O.E., and C.S. wrote the manuscript with contributions from all authors.

## ACKNOWLEDGEMENTS

- This work was supported by the National Institutes of Health (K08-AI089938 to CS) University of Washington Department of Medicine and UW Royalty Research Fund (CS).
- The authors would like to thank Branch Moody and Martine Gilleron for providing lipid antigens and John Hansen and the Research Cell Bank at the Fred Hutchinson Cancer Research Center for assembling the cohort of bone marrow donors. We acknowledge the NIH Tetramer Core Facility (contract HHSN272201300006C) for provision of biotinylated CD1B monomers.

